# Genotypic similarity among algal symbionts corresponds to associations with closely related coral hosts

**DOI:** 10.1101/2020.09.26.314773

**Authors:** Hannah G. Reich, Sheila A. Kitchen, Kathryn H. Stankiewicz, Meghann Devlin-Durante, Nicole D. Fogarty, Iliana B. Baums

## Abstract

Mutualisms where hosts are coupled metabolically to their symbionts often exhibit high partner fidelity. Most reef-building corals form obligate symbioses with specific species of photosymbionts, dinoflagellates in the family Symbiodiniaceae, despite needing to acquire symbionts early in their development from environmental sources. Three Caribbean acroporids (*Acropora palmata*, *A. cervicornis*, and their hybrid *A. prolifera*) are geographically sympatric across much of their range in the greater Caribbean, but often occupy different depth and light habitats. Both species and their hybrid associate with *Symbiodinium ‘fitti’*, a genetically diverse species of symbiont that is specific to these hosts. Since the physiology of the dinoflagellate partner is strongly influenced by light (and therefore depth), we investigated whether *S. ‘fitti’* populations from each host source were differentiated genetically. We generated shallow genome sequences of acroporid colonies sampled from across the Caribbean. Single Nucleotide Polymorphisms (SNPs) among *S. ‘fitti’* strains were identified by aligning sequences to a ~600 Mb draft assembly of the *S. ‘fitti’* genome, assembled from an *A. cervicornis* metagenome. Phylogenomic and multivariate analyses revealed that allelic variation among *S. ‘fitti’* partitioned to each host species, as well as their hybrid, rather than by biogeographic origin. This is particularly noteworthy because the hybrid, *A. prolifera*, has a sparse fossil record and may be of relatively recent origin. Many of the SNPs putatively under selection were non-synonymous mutations predicted to alter protein efficiency. Differences in allele frequency among *S*. ‘*fitti*’ populations from each host taxon may correspond to distinct phenotypes that thrive in the different cellular environments found in each acroporid. The non-random sorting among genetically diverse strains, or genotypes, to different hosts could be the basis for lineage diversification via disruptive selection, leading to ecological specialization and ultimately speciation.

## Introduction

Ecosystem services provided by coral-dinoflagellate mutualisms rival the contributions of other widely studied symbioses (ex: tubeworms and bacteria, yeast and termites, plant and fungi). In the coral-dinoflagellate mutualism, each partner benefits as the coral receives photosynthetic sugars from their dinoflagellate symbionts and the algal symbiont receives nutrients and protection in return (Trench 1979). Though decades of investigations have probed the causes and consequences of coral-algal dysbiosis (ex: coral bleaching), we are still gathering information on their basic biology (Cziesielski, Schmidt-Roach, & Aranda 2019). Specifically, little is known about the intraspecies level of co-evolutionary dynamics between host and their symbionts. The interdependence of the partners adds complexity to the system as each partner is selected in the context of the other.

Understanding the mechanisms that promote the apparent fidelity of reef-building corals towards one endosymbiotic dinoflagellate species (Symbiodiniaceae), despite having the opportunity to horizontally acquire symbionts, is important in light of rapid climate change. This is because the ability of corals to cope with heat stress by “shuffling” their endosymbiont communities to a more heat tolerant lineage (Little, Oppen, & Willis 2004; Silverstein, Cunning, & Baker 2017) may be limited. Corals have co-evolved with the Symbiodiniaceae since the Jurassic period (LaJeunesse et al. 2018) and, over time, may have become uniquely adapted to their symbionts (Goulet 2006; LaJeunesse et al. 2018; LaJeunesse et al. 2004; Lewis, Chan, & LaJeunesse 2019; Parkinson, Coffroth, & LaJeunesse 2015b; Thornhill, Fitt, & Schmidt 2006a). While juvenile corals often host several symbiont species, this community wanes over time to the dominant symbiont species (Abrego, Van Oppen, & Willis 2009; Coffroth, Goulet, & Santos 2001; Poland & Coffroth 2017). This suggests that corals are most compatible with their dominant symbiont species and foreign pairings might be maladaptive, at least under current conditions (Cunning, Gillette, Capo, Galvez, & Baker 2015; Pettay, Wham, Smith, Iglesias-Prieto, & LaJeunesse 2015).

A central question in coral reef science is whether coral-dinoflagellate symbioses can adapt to increasing sea surface temperatures over ecological time scales, especially if shuffling of symbiont partners is restricted (Goulet 2006). Genetic variation present within algal species has remained largely unstudied as a source that may fuel such adaptation (Buerger et al. 2020; Parkinson, Banaszak, Altman, LaJeunesse, & Baums 2015a). The potential for genetic variation within algal species to fuel adaptation to changing conditions can be assessed in the laboratory via experimental evolution experiments where algal strains are selected over several generations under heat stress conditions (Baker et al. 2018; Buerger et al. 2020; Chakravarti & van Oppen 2018). In one instance, Symbiodiniaceae strains adapted to heat stress selection *in vitro* but once introduced into the coral partner, gains were not always retained highlighting the complexity of adaptation in the context of mutualistic partners (Buerger et al. 2020).

Alternatively, field studies may assess the long-term influence of selective factors such as a strong light gradient on the genetic variation of Symbiodiniaceae by taking advantage of depth stratification found in their coral hosts (Bongaerts et al. 2015a; Serrano et al. 2016). While sharing a geographic range, Caribbean *Acropora* species often differentiate across a depth and light gradient; *A. cervicornis* occupies a lower light habitat (~10 m depth) relative to its high-light dwelling (~3 m depth) sibling species *A. palmata* and their hybrid *A. prolifera* (~1 m depth; (Fogarty 2012; Goreau 1959; LaJeunesse 2002). All three taxa harbor the dinoflagellate endosymbiont *Symbiodinium ‘fitti’* (ITS2 type A3), which is distinct from other *Symbiodinium* A3 lineages found in giant clams and other cnidarians (Kemp et al. 2015; Lee et al. 2015; Pinzón et al. 2015; Shoguchi et al. 2018). The variation of morphology between the three taxa ranges from broad, moose antler branches to thinner, stag antler branches results in differences in the flow and light field within and around the colonies (Enríquez, Méndez, Hoegh-Guldberg, & Iglesias-Prieto 2017; Gladfelter 1983; Gladfelter 2007). Therefore, the persistence of *S. ‘fitti’* in three host taxa at a range of depths across a large geographic region provides a unique opportunity to study how evolutionary history, geography, natural ecology, and biophysical parameters may contribute to adaptation and co-evolution in the coral holobiont.

The ecological and evolutionary dynamics between host and symbiont species are influenced by differences in their reproduction and dispersal strategies (Reviewed in Thornhill, Howells, Wham, Steury, & Santos 2017). Caribbean acroporid corals reproduce via production of meiotic, planktonic larvae and also disperse locally via fragmentation. Gene flow is restricted between eastern and western Caribbean region for *A. palmata* and *A. cervicornis* (Baums, Miller, & Hellberg 2005, 2006). Within each region, further population structure is observed but the specifics differ between species with *A. cervicornis* showing generally more fine-scale differentiation then *A. palmata* (Baums et al. 2005, 2006; Devlin-Durante & Baums 2017; Hemond & Vollmer 2010; Kitchen et al. 2019; Vollmer & Palumbi 2002, 2006). *A. palmata* and *A. cervicornis* have been present in the fossil record since the late Pliocene (~2.6-3.6 Mya) whereas the hybrid *A. prolifera* is mostly absent from the fossil record (Budd & Johnson 1999; McNeill, Budd, & Borne 1997; Precht, Vollmer, Modys, & Kaufman 2019). Although *A. prolifera* produces viable eggs and sperm, molecular analyses indicates that F2 adults are very rare or absent while backcrosses with either parent species occur occassionally (Kitchen et al. 2020; Van Oppen, Willis, Vugt, & Miller 2000; Vollmer & Palumbi 2002).

The population structure and genotypic diversity of *S. ‘fitti’* has received less attention and at a coarser level of genomic resolution (Baums et al. 2019; Baums, Devlin-Durante, & LaJeunesse 2014; Thornhill et al. 2017). Despite that, higher levels of population genetic structure are documented in *S. ‘fitti’* when compared to one of its hosts, *A. palmata* (Baums et al. 2014). *S. ‘fitti’* cells divide mitotically within the host and water column dispersal appears limited (Fitt & Trench 1983; Thornhill et al. 2017). Accordingly, the majority of Caribbean acroporids colonies host a single strain of *S. ‘fitti’* and maintain long-term fidelity to that strain (Baums et al. 2014; O’Donnell, Lohr, Bartels, Baums, & Patterson 2018). However, sexual reproduction in Symbiodiniaceae has not been completely ruled out as a reproductive strategy because meiotic machinery has been detected in genomic data (Bellantuono, Dougan, Granados-Cifuentes, & Rodriguez-Lanetty 2019; Chi, Parrow, & Dunthorn 2014; Levin et al. 2016; Shah, Chen, Bhattacharya, & Chan 2020) and recombination is evident in population genetic data (Baums et al. 2014). These contrasting life-history strategies may contribute to the higher levels of population structure of *S. ‘fitti’* compared to their acroporid hosts throughout the Caribbean (Baums et al. 2014; Thornhill et al. 2017).

Here, we describe fine-scale genetic differences in *S. ‘fitti’* strains across its three host taxa spanning the geographic distribution of the mutualism. A draft *S. ‘fitti’* genome assembly was constructed from *A. cervicornis* metagenomic sequences and compared to other Symbiodiniaceae genomic resources. Variation in genome-wide single nucleotide polymorphisms (SNPs) were investigated in *S. ‘fitti’* and scanned for mutations that may change protein structure and function. Lastly, the potential biological and evolutionary ramifications of the allelic composition of *S. ‘fitti’* are discussed.

## Methods

### Sample Collection, Sequencing, and Assembly

Tissue was collected for genome sequencing from 76 acroporids spanning the geographic distribution of *S. ‘fitti’* (Fig. 1, Table S1; Kitchen et al. 2019). High molecular weight DNA was isolated from each coral tissue sample using the Qiagen DNeasy kit (Qiagen, Valencia, CA) without prior enrichment for *S. ‘fitti’*. Of these samples, one specimen for each species from Florida (*A. cervicornis* CFL14120 and *A. palmata* PFL1012) was ‘deeply’ sequenced (~150x coverage; Kitchen et al. 2019). Paired-end short insert (550 nt) sequencing libraries of the two deeply sequenced samples were constructed with 1.8-2 μg sample DNA and the TruSeq DNA PCR-Free kit (Illumina, San Diego, CA). The remaining 74 paired-end short insert (350 nt) sequencing libraries were constructed using 100 ng sample DNA and the TruSeq DNA Nano kit (Illumina, San Diego, CA) with coverage between 8-40x (Kitchen et al. 2019). Deep- and shallow-sequence libraries were pooled separately and sequenced on either Illumina HiSeq 2500 or HiSeq 4000 (Table S1, Illumina, San Diego, CA).

**Fig. 1:**
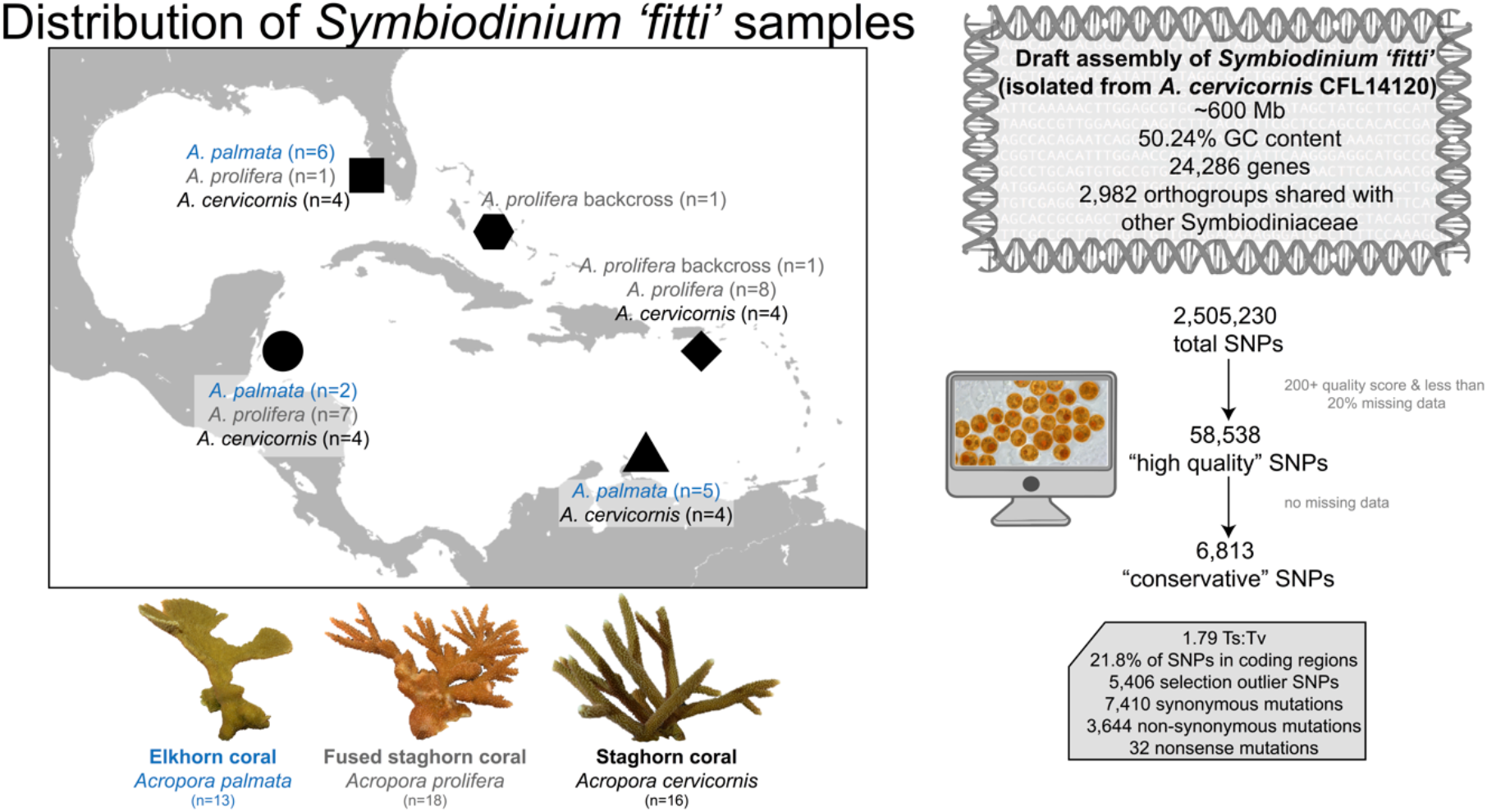
Sampling design and summary statistics of population genomic approach used to characterize *Symbiodinium ‘fitti’* across three coral hosts. *Acropora palmata* (n = 13)*, Acropora cervicornis* (n = 16), and *Acropora prolifera* (n = 18) samples used for shallow genome sequencing spanned their geographic distribution. Summary statistics for the ‘deeply-sequenced’ CFL 14120 *A. cervicornis – S. ‘fitti’* draft genome assembly including overall length, %GC content, # of genes, and shared gene families with other Symbiodiniaceae. Summary statistics and visual depiction of quality filtering work flow that was employed to identify high quality variants and those that are under selection. Coral images from N. Fogarty and I. Baums.

Sequencing adaptors and low-quality base calls (Phred score < 25) from the 3’ end of the deeply sequenced *A. cervicornis* metagenome reads were trimmed using cutadapt v 1.6 (Martin 2011). After initial filtering, processed reads shorter than 50 bp were discarded and PCR duplicates removed using FastUniq v. 1.1 (Xu et al. 2012). A series of filtering steps were completed to identify the fraction of reads originating from *A. cervicornis* and *Symbiodinium* spp. First, a modified approach similar to Blobology, which compares sequence homology, read coverage and GC content, was performed (Kumar, Jones, Koutsovoulos, Clarke, & Blaxter 2013). Contigs from a preliminary genome assembly created with SOAPdenovo2 v0.4 (parameters −K 95 –R) were compared for homology against the coral *Acropora digitifera* (NCBI: GCF_000222465.1) and symbiont *Breviolum minutum* (OIST: symbB.v1.0.genome.fa) genomes, and NCBI nucleotide database nt using megablast (evalue 1e-5; Altschul et al. 1997; Shinzato et al. 2011; Shoguchi et al. 2013). The sequence matches to the nt database for contigs that had no match to either coral or symbiont genome, were used to create a local contamination database to further screen the reads (Luo et al. 2015).

Reads were aligned with Bowtie2 v. 2.2.9 (parameters –q –fast; Langmead & Salzberg 2012) consecutively to the *A. digitifera* mitochondria (KF448535.1), followed by a concatenated set of three Symbiodiniaceae genomes (*Symbiodinium microadriaticum, Breviolum minutum, Fugacium;* Aranda et al. 2016; Lin et al. 2015; Shoguchi et al. 2013), and the contamination database. This filtering step, however, only aligned 0.28% of the reads from *A. cervicornis* to the Symbiodiniaceae genomes. Reads that mapped to Symbiodiniaceae genomes (n=1,004,992) were extracted and assembled using SPADES v3.9.1 with a multi-kmer approach (−k 21,33,55,77,99) (Bankevich et al. 2012). The reads that aligned to the contamination database were assembled separately with SPADES v3.9.1 as described above, resulting in three additional contigs that matched Symbiodiniaceae genomes through blast homology.

The filtered *A. cervicornis* reads were assembled with SoapDeNovo v2, followed by six rounds of gap filling using GapCloser v1.12 and scaffolding using both SSPACE v2.0 (rounds 1, 3, and 5) and LINKs v1.8.5 with *A. digitifera* scaffolds as the “long-reads” (−t 2 –d 3000 –k 25, round 2, 4 and 6) on alternate rounds (Boetzer & Pirovano 2014; Luo et al. 2015; Warren et al. 2015). After the first three rounds and then each subsequent round, the contigs/scaffolds were partitioned to either coral or symbiont based on the top scoring match (lowest e-value) against a local blast database containing three Symbiodiniaceae genomes and five cnidarian genomes (*Hydra*, *Hydractinia*, *Nematostella, Exaiptasia, Acropora digitifera*). If the scaffolds equally matched cnidarian and symbiont sequences or did not match either they were retained in the coral fraction. Scaffolds identified as Symbiodiniaceae after the six rounds as well as those assembled with SPADES above were combined. Two additional rounds of scaffolding with LINKS using *S. microadriaticum* as “long-reads” followed by SSPACE and gap filling with GapCloser were performed. To remove any remaining sequences matching cnidarian sequences, a final round of scaffold partitioning was performed by comparing the scaffolds against the *Acropora* spp. genomes: *A. digitifera* (Shinzato et al. 2011), *A. palmata* (Kitchen et al. unpublished; *http://baumslab.org/research/data), A. cervicornis* (Kitchen et al. unpublished)*, A. hyacinthus* (Barshis et al. 2013)*, A. tenuis* (Liew, Aranda, & Voolstra 2016); and *Symbiodinium* spp. genomes: *S. microadriaticum* (Aranda et al. 2016), and *S. tridacnidorum* (Shoguchi et al. 2018).

### Genome annotation and completeness

Genes were predicted using Augustus v 3.2.3 (Stanke et al. 2006; Stanke, Steinkamp, Waack, & Morgenstern 2004). Each predicted gene in *S. ‘fitt*i’ was queried against the NCBI nr, Uniprot SwissProt and trembl databases using blastx 2.6.0+ (max target seqs = 5, max hsps = 1, e-value = 1e-5; Altschul et al. 1997; Apweiler et al. 2004; Bairoch & Apweiler 1997; UniProt 2014). Gene models were also compared to the *S. microadriaticum* gene and protein predictions (NCBI: GCA_001939145.1) using blast (Altschul et al. 1997). To calculate assembly statistics and compare completeness to other Symbiodiniaceae genomes available at the time of analysis (*F. kawagutii, C. goreaui, B. minutum, S. microadriaticum, S. tridacnidorum*), we compared our *S. ‘fitti’* assembly and the aforementioned assemblies using an online version of CEGMA with the eukaryote ortholog set executed by gVolante (https://gvolante.riken.jp; Simão, Waterhouse, Ioannidis, Kriventseva, & Zdobnov 2015). Orthofinder v2.2.1 with default settings (Emms & Kelly 2015) was used to identify unique and shared orthogroups between *S. ‘fitti’* and six other Symbiodiniaceae species.

### S. ‘fitti’ infection status

Presence of multiple *S. ‘fitti’* strains within a coral host sample was determined using 12 *S. ‘fitti’* specific microsatellite loci as described by Baums et al. (2014). *S. ‘fitti’* is haploid, thus samples with multiple alleles for any given *S. ‘fitti’* microsatellite locus were deemed co-infected and removed from downstream analyses (Table S1).

### Variant detection and filtering

For the purpose of SNP analyses, the *S. ‘fitti’* genome assembly based on the *A. cervicornis* metagenome was used as a reference for variant calling of the deeply-sequenced *A. palmata* and all shallow genome samples (Table S1). The 47 shallow *S. ‘fitti’* and 1 deep sequenced *A. palmata* “like” *S. ‘fitti’* genome samples were aligned using BWA v0.7.15 (Li 2013). Samtools v1.4 was used to remove PCR duplicates from the BAM file and alignment statistics were calculated using samtools *flagstat* (Table S1; Li et al. 2009). Variants were gathered using samtools *mpileup* using the –ugAEf and –t AD,DP flags and called using bcftools v1.4 using the haploid, –f GQ, and -vmO z flags (Li et al. 2009; Narasimhan et al. 2016). The bcftools (Li et al. 2009; Narasimhan et al. 2016) –m2 –M2 –v snps flags were used to separate SNPs from the output and the –v indels flag was used to remove indels from the output (Narasimhan et al. 2016). High-quality SNPs and indels were characterized as variants with a quality score over 200 and with no more than 20% of variant calls missing at a given site among all samples (Danecek et al. 2011; Narasimhan et al. 2016). Only high-quality SNPs were used in subsequent analyses.

### Population structure

The *psbA* minicircle was assembled from each sample to determine if the dominant algal partners amongst the three host taxa were all *S. ‘fitti’* The psbA minicircle in the *S. ‘fitti’* genome assembly was identified through blast searches against three psbA sequences from NCBI (JN557866.1 = *Symbiodinium* type A3, JX094319.1 = *Breviolum minutum*, and AJ884898.1= *B. faviinorum* (Barbrook, Visram, Douglas, & Howe 2006; Mungpakdee et al. 2014; Pochon, Putnam, Burki, & Gates 2012). The *psbA* minicircle for the remaining samples was assembled using two approaches. In the first approach, filtered and trimmed short-read sequences were mapped to *S. ‘fitti’ psbA* sequence (scaffold71443) using Bowtie 2 v2.3.4.1 (Langmead & Salzberg 2012) with the --sensitive mode parameter. Mapped reads were extracted using bedtools v2.26.0 (Quinlan & Hall 2010) and assembled using SPAdes v3.10.1 (Bankevich et al. 2012) with various kmer sizes (−k 21, 33, 55, 77 and 99). In the second approach, the *de novo* organelle genome assembler NOVOplasty was used (Dierckxsens, Mardulyn, & Smits 2016). The *S. ‘fitti’* genome *psbA* sequence was used as the seed sequence to extract similar sequences from the original, unfiltered reads for each sample. A consensus sequence from the two approaches for each sample was created after manual alignment of the sequences using MEGA6 (Tamura, Stecher, Peterson, Filipski, & Kumar 2013).

Phylogenomic patterns of *S. ‘fitti’* allelic composition were determined using a subset of the high-quality SNPs without missing data with the RAxML-NG v. 0.9.0 GTR+FO+G nucleotide model (Stamatakis 2014). The tree topology with the lowest likelihood score is presented with nodal support from 100 bootstrap replicates (Stamatakis 2014). Population structure was evaluated using STRUCTURE v2.3.4 for the 58,813 high-quality SNPs (Pritchard, Stephens, & Donnelly 2000). Additionally, the R package poppr v2.1.0 was used to determine the multilocus genotype of each strain using high quality SNPs with different genetic distance thresholds ranging 10-20% (See Table S1; Kamvar, Tabima, & Grünwald 2014; Kitchen et al. 2020) Clusters in multivariate space were detected using the *pca* function in PCAdapt (Luu, Bazin, & Blum 2017). An Analysis of Molecular Variance (AMOVA, poppr R package) was used for additional detection of population differentiation (Kamvar et al. 2014).

### Determination of variants under selection

Two different methods were used to identify candidate loci under selection. BayeScan v2.1 is a Bayesian method that incorporates uncertainty of allele frequencies between populations with small sample sizes (Fischer, Foll, Excoffier, & Heckel 2011; Foll, Fischer, Heckel, & Excoffier 2010; Foll & Gaggiotti 2008). The default BayeScan settings were used for determining SNPs under selection when accounting for *S. ‘fitti’* host, location, and host*location interactions. PCAdapt v4.0.3 was used to determine SNPs under selection without prior population information using the default settings (Knaus & Grünwald 2017). Outliers from BayeScan were determined as markers where FDR <0.05 and outliers from PCAdapt v4.0.3 were determined by q-values larger than the alpha value (0.05; Fischer et al. 2011; Foll et al. 2010; Foll & Gaggiotti 2008; Knaus & Grünwald 2017). Outlier loci with a Bayes probability of 1 and q-value of 0 which becomes infinite following logarithmic transformation and were removed from plotting. All statistics from SNPs under selection, their proximity to coding regions, sequence coverage, and per SNP F_ST_ are in Tables S7 and S9. SnpEff v4.3 was used to predict downstream functional implications of all detected variants (De Baets et al. 2011).

### Data and code availability

Raw data is publicly available on NCBI under SRA project PRJNA473816. Code for data analysis and figure generation is available on github (https://github.com/hgreich/Sfitti).

## Results

### Genome statistics and comparison to other Symbiodiniaceae

The *Symbiodinium ‘fitti’* assembly has a total nucleotide length of over 600 Mb (601,782,011 bp) and contains 274,185 contigs/scaffolds (*A. cervicornis-S. ‘fitti’* CFL14120; Fig. 1, Table S2). The *A. palmata-S. ‘fitti’* (PFL1012) deeply-sequenced sample had 297,371,995 reads map to the *A. cervicornis-S. ‘fitti’* (CFL14120) reference (19% mapping rate, 8.5% paired reads, 4.2% singleton reads; Table S1). The shallow-sequenced genome samples with one *S. ‘fitti’* genotype had an average of 6,461,332 reads map to the reference (19.2% mapping rate, 8.8% paired reads, 4.3% singleton reads; Table S1). The GC content and number of ambiguous bases were comparable to the *S. microadriaticum* assembly at 50.24% and 4.82%, respectively (Fig. 1, Table S2; Aranda et al. 2016). The genome completeness was assessed by the identification of the 248 core genes queried using the CEGMA program. The *S. ‘fitti’* assembly had 55 complete proteins, 79 complete + partial proteins, and 169 missing proteins, which is comparable to the other Symbiodiniaceae assemblies queried (Table S2). The average number of orthologs per core gene was ~1.4 for the Symbiodiniaceae assemblies (Table S2). Gene prediction of *S. ‘fitti’* assembly revealed 24,286 gene models, however, many were incomplete (i.e., missing start or stop codon; Table S2, S3). In the gene family analysis, 3,368 orthogroups were found to be shared by the Symbiodiniaceae assemblies excluding *S. ‘fitti’* whereas 2,982 orthogroups were found to be shared by all Symbiodiniaceae assemblies (Fig. S1; including *S. ‘fitti’*). Additionally, 1,898 orthogroups were uniquely shared by *S. ‘fitti’* and its closest relative *S. tridacnidorum* whereas 1,357 orthogroups were shared by the three assemblies from the genus *Symbiodinium* (Fig. S1). *S. ‘fitti’* had 11 orthogroups that were not shared with any other Symbiodiniaceae assemblies.

### Variation of the allelic composition of Symbiodinium ‘fitti’

Based on the analysis of *S. ‘fitti’* specific microsatellite loci, the majority of shallow-sequenced samples harbored one strain of *S. ‘fitti* (n= 47, 75.9% of *A. palmata-S. ‘fitti’* and 82.6% of *A. cervicornis-* and *A. prolifera-S. ‘fitti’*; Table S1) and were used for further analysis. A total of 2,505,230 SNPs and 569,337 indels were identified between all samples. Of these, 58,538 SNPs and 1,874 indels were considered high-quality (Fig. 1). The 58,538 high-quality SNPs represent a range of per SNP fixation levels from 0 to 1 (Fig. S2). The Transition: Transversion ratio of the high-quality SNPs was 1.79 and did not vary by host species (Table S1). Of the high-quality SNPs, 12,780 (21.8%) occurred in coding regions (Table S4). The majority of the SNPs (87.5%) in coding regions matched other Symbiodiniaceae genomic resources (primarily the closely related species, *S. microadriaticum*). Multi-locus genotype (MLG) filtering of the 58,538 “high quality” SNPs indicated each sample represented a unique MLG (strain), consistent with the microsatellite analysis, and was retained for downstream analyses (Table S1). Additional filtering to remove variants with missing data resulted in 6,813 high-quality SNPs, hereafter called conservative SNPs. After this procedure of quality filtering SNPs and setting a stringent missing data threshold, the average read coverage increased from 1.53 to 11.3 per SNP (Table S1; average 655.3% increase).

### Patterns of host-specificity and biogeography within S. ‘fitti’

The phylogeny of the *psbA* minicircle non-coding region revealed little differentiation between symbiont strains with respect to their host taxa, confirming that *S. ‘fitti’* is one species (Fig. S2). The AMOVA corroborated that most of the variation among *S. ‘fitti’* is at the within-species level (86.1+%; Table S5). Variation attributed to host species explained 11.6% of the components of covariance (sigma = 87.0) and then variation among the geographic location of each host explained 2.3% of the components of covariance (sigma = 17.1; Table S5). Variation among geographic locations explained a negative amount of the components of covariance (−5.4%, sigma = −39.3) whereas variation among the host taxa at the various geographic locations explained 16.3% of the components of covariance (sigma = 117.6; Table S5).

Consistent with the AMOVA results, samples clustered primarily by host taxon rather than geographic origins in a Principal Component Analysis (PCA) with the high-quality SNPs and in the Maximum Likelihood tree with the conservative SNPs (Figs. 2, 3). In the PCA, 16% of variance was explained by PC1 whereas 13.6% was explained by PC2 (Fig. 2). Within each host, there was some indication of biogeographic partitioning in the phylogeny but not in the PCA (Figs. 2, 3; Table S6). The *S. ‘fitti’* associated with *A. prolifera* were found intermediate to the parental species in the Maximum Likelihood tree and clustered loosely together but were more similar to the *A. palmata S. ‘fitti’* (Fig. 2). Analysis of STRUCTURE output using the *delta K* method (Evanno, Regnaut, & Goudet 2005) identified three clusters as the most likely K (Table S6). The three clusters largely corresponded to host taxa (Fig. 4).

**Fig. 2:**
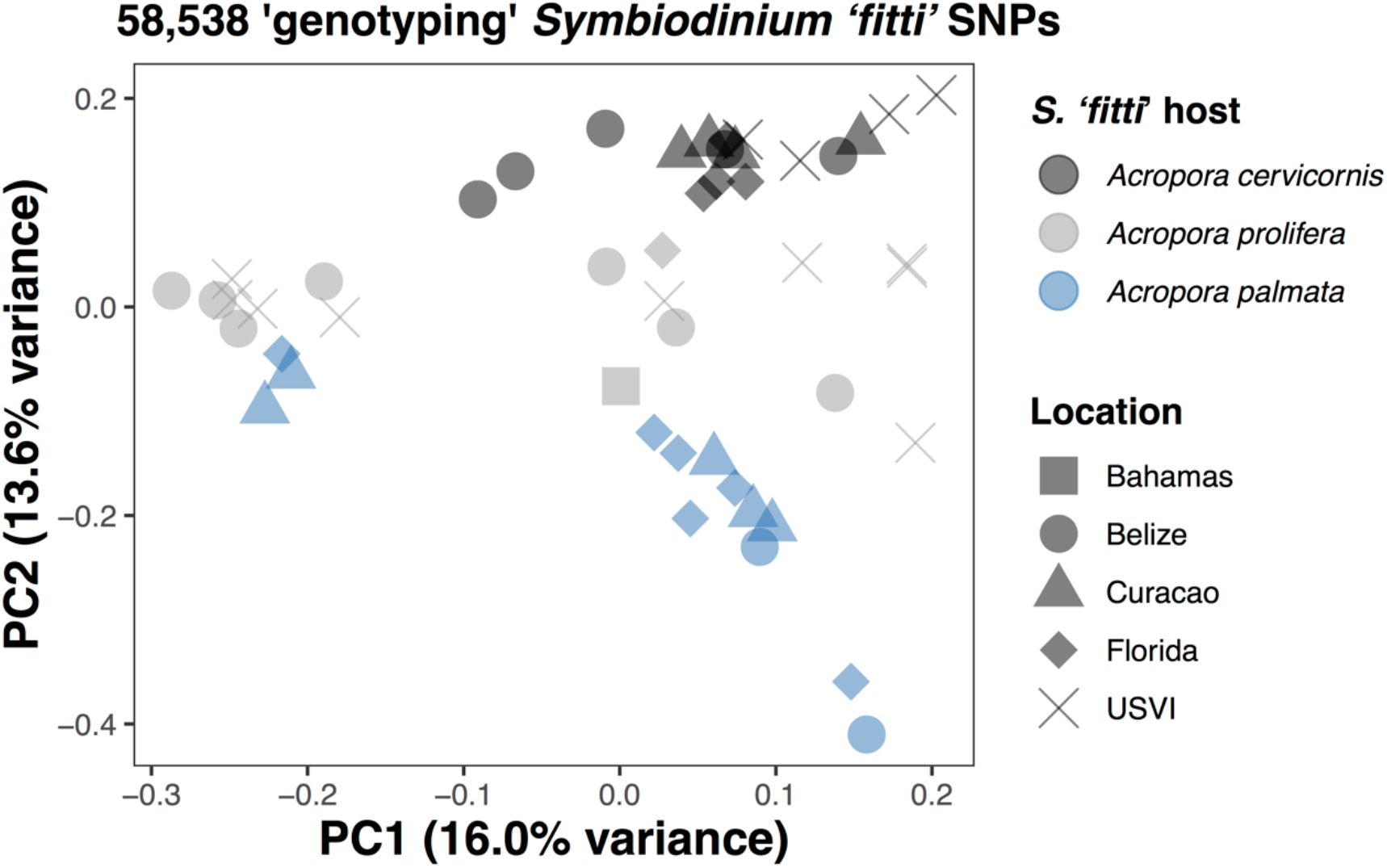
Principal component analysis (PCA) of 58,583 genotyping *Symbiodinium ‘fitti’* SNPs illustrates genomic differentiation by host taxon. Coral images from N. Fogarty and I. Baums.

**Fig. 3:**
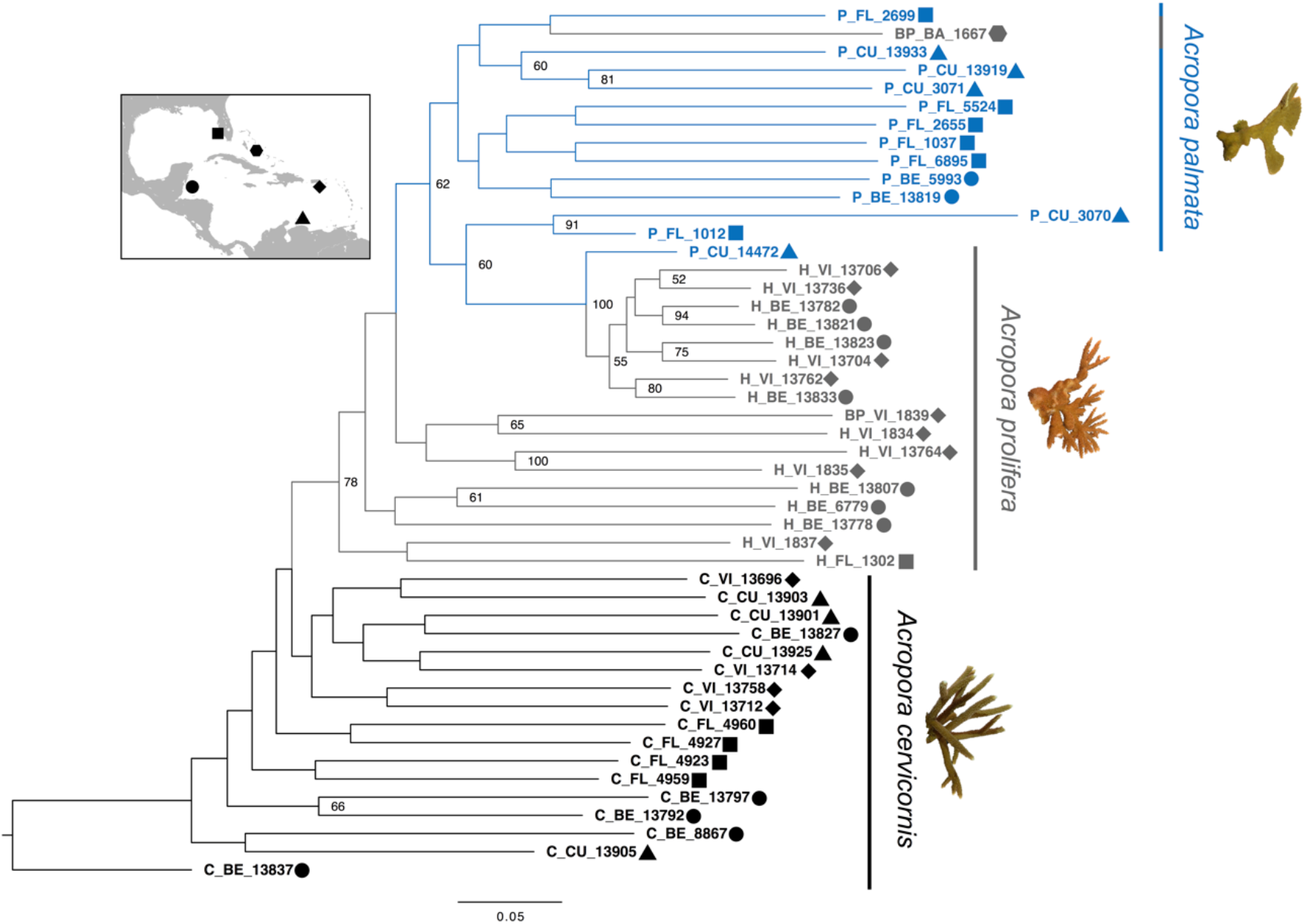
Allelic composition of *Symbiodinium ‘fitti’* is at the sub-species level. RAxML (Maximum Likelihood) phylogeny of 6,813 “conservative” genotyping *S. ‘fitti’* SNPs without missing data and 100 bootstrap replicates illustrate *S. ‘fitti’* genomic differentiation by host taxon.

**Fig. 4:**
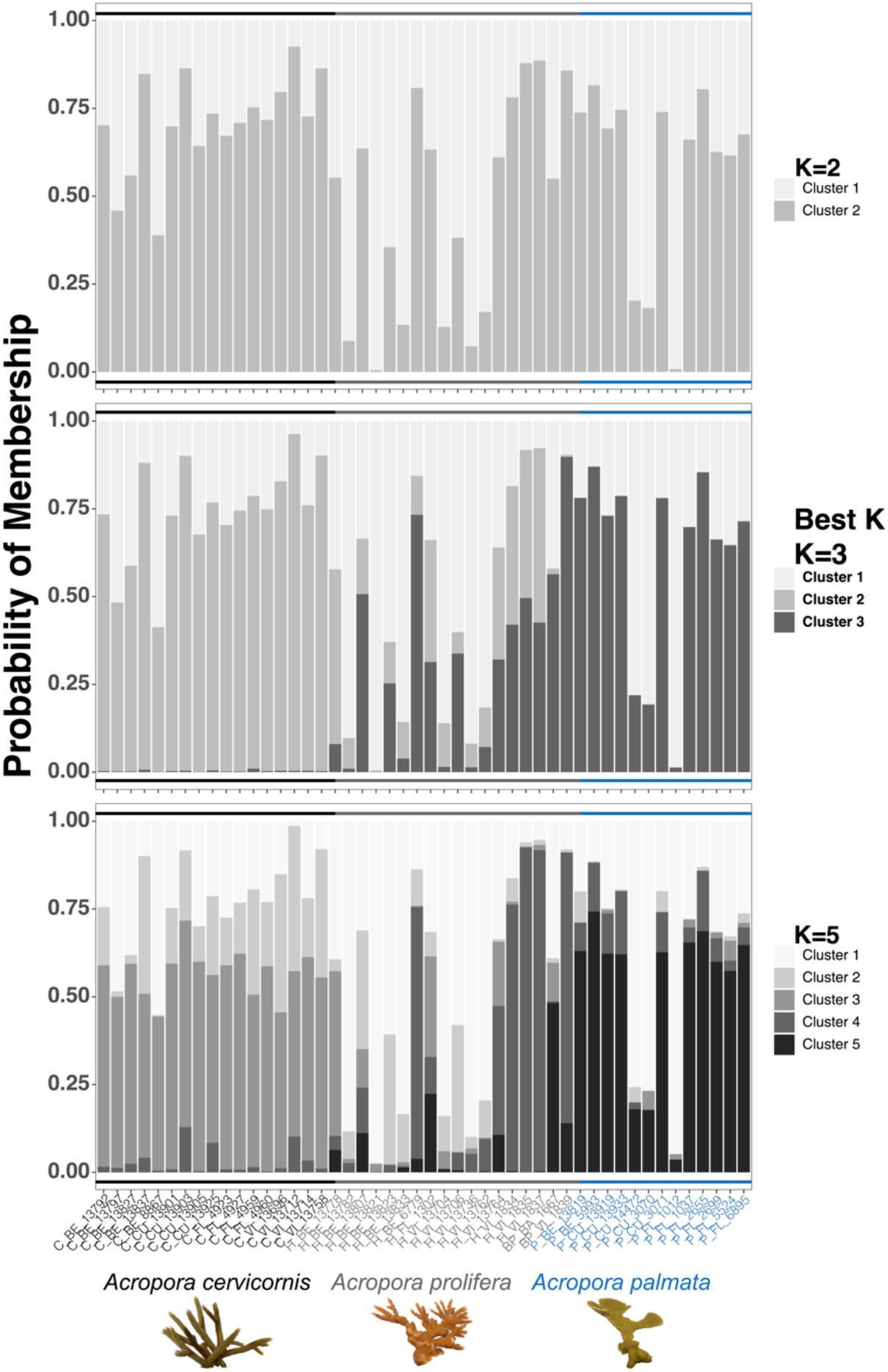
*Symbiodinium ‘fitti’* strain population assignment aligns with host taxon. Probability of membership predicted by STRUCTURE for 58,583 “high quality” *Symbiodinium ‘fitti’* SNPS. K=3 was determined as best K. K=2 and K=5 are also presented for comparison. These results illustrate that *S. ‘fitti’* membership clusters largely correspond to their host acroporid taxon. Coral images from N. Fogarty and I. Baums.

### SNPs under selection

Of the high-quality SNPs, 4,987 (8.5%) were determined as selection outliers by PCAdapt (Fig. 5). When BayeScan accounted for host identity, location of host, and host by location interaction, 217, 5 and 197 selection outliers were identified, respectively (n= 370 SNPs; Fig. 5; Table S7). Additionally, 339 selection outlier SNPs were shared between the two programs (Fig. 5; Table S7). 103 outlier loci identified by BayeScan had a Bayes probability of 1 and q-value of 0 which becomes infinite following logarithmic transformation and were therefore removed from the Manhattan plot (Fig. 5) but were reported in Table S7. For each set of SNPs under selection, a subset was found in coding regions (899 from PCAdapt, 19 from BayeScan, 14 from both callers; Table S7). The 14 outliers in coding regions that were shared by both callers were found in the coding regions of Putative cytosolic oligopeptidase, tankyrase-like proteins, alpha-agarase, and uncharacterized proteins (Table S7).

**Fig. 5:**
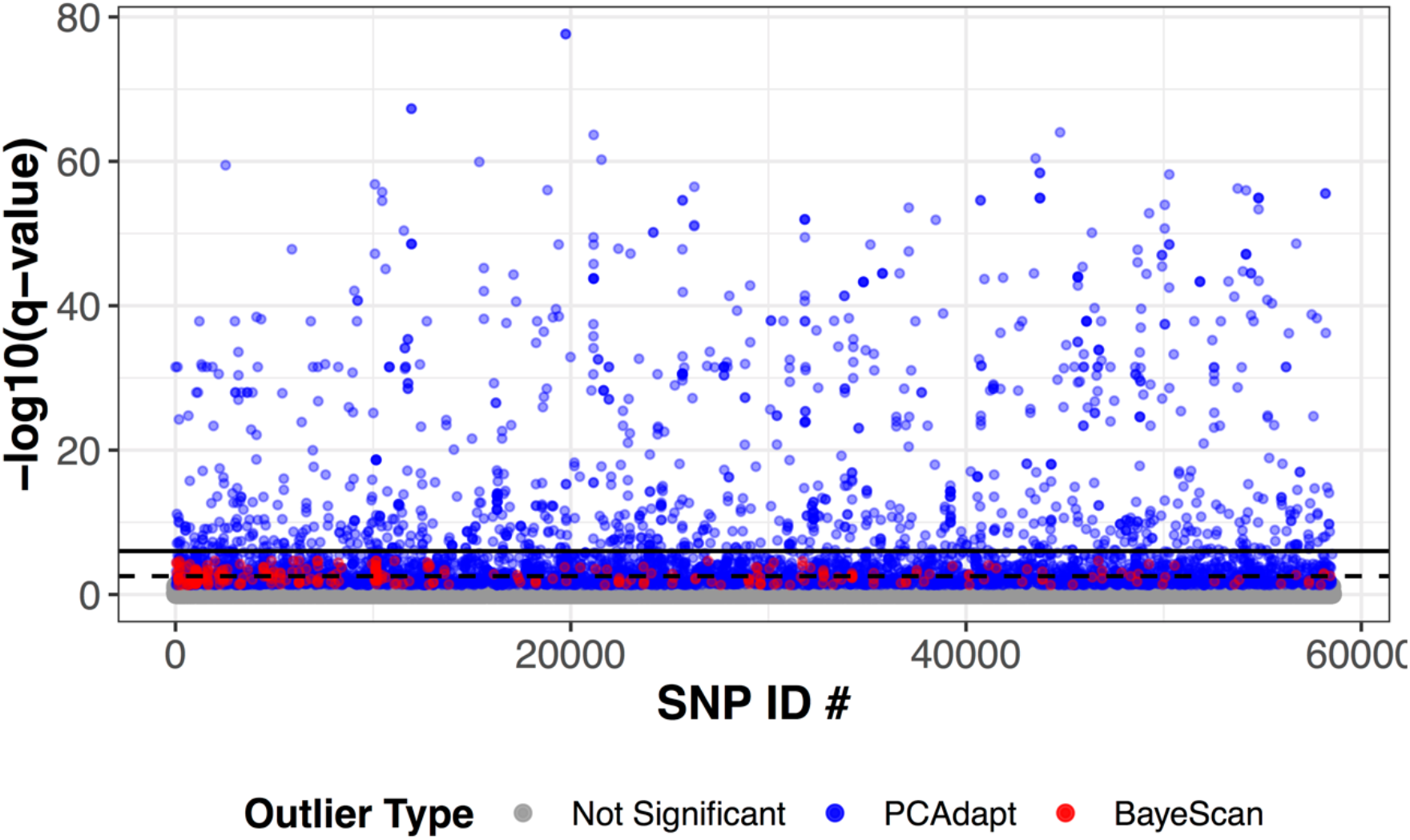
Genetic variants of *S. ‘fitti’* showing signatures of selection. Manhattan plot of –log10 transformed q-values for 58,583 genotyping SNPs. SNPs are highlighted by outlier detection program (PCAdapt loci highlighted in blue; BayeScan loci highlighted in red). 339 selection outlier SNPs were shared between the two programs. 103 outlier loci identified by BayeScan had a Bayes probability of 1 and q-value of 0 which becomes infinite following logarithmic transformation and were therefore removed from the plot. These loci were also had high BayeScan fixation levels between host and location. The dashed line represents the 5% FDR adjustment threshold (2.54) and whereas the solid line represents the 0.05 Bonferroni correction threshold (6.01).

### Predicted functional implications of SNPs within S. ‘fitti’

Of all high-quality SNPs, SnpEff identified 60,373 modifier/non-coding variants (84.43%), 3,629 moderate/mostly harmless variants (5.08%), 7,451 low impact variants that might change protein efficiency/effectiveness (10.42%), and 51 highly disruptive SNPs (0.07%). Of the predicted mutations, SnpEff identified 3,644 non-synonymous mutations (32.87%), 32 premature stop codon/nonsense mutations (0.29%), and 7,410 synonymous mutations (66.84%; Table S8). The aforementioned mutations are predicted to cause 11,086 codon changes and 3,676 amino acid changes (Tables S8, S9). SnpEff predicted the 14 aforementioned outlier mutations of Putative cytosolic oligopeptidase, tankyrase-like proteins, and alpha-agarase as modifier variants found in the introns (Tables S7-S10).

## Discussion

The population dynamics and evolutionary history of reef-building corals are relatively well studied compared to the dinoflagellate symbionts they harbor. However, selection acts on both partners and differences in life-history characteristics between algae and corals suggest that the spatial and temporal scale of adaptation may differ. This would have consequences for our understanding of how they may adapt to rapidly changing climates. Genomic data of Caribbean acroporids reveals fine-scale population structure within the host taxa (Devlin-Durante & Baums 2017; Kitchen et al. 2019; Kitchen et al. 2020). Previous analyses of microsatellite loci demonstrated that *S. ‘fitti’* gene-flow scales were smaller than its host *Acropora palmata* (Baums et al. 2014), however, this study jointly analyzed SNP data from all three Caribbean acroporid host taxa. We showed that the allelic composition of sympatric *S. ‘fitti’* populations are partitioned by host taxon and describe two potential scenarios that would lead to this result (Figs. 2–4). Differentiation of *S. ‘fitti’* by host taxon implies partner selectivity, which may be the result of coevolution (Scenario 1). Alternatively, coevolution and partner selectivity *per se* (i.e. via specific recognition of genetic variants among partners) may not explain the patterns of allelic variation. Differences in the micro-environment (light, depth, nutrient availability) associated with the habitat preferences of the host taxa may drive symbiont differentiation without specific recognition interactions or coevolution (Scenario 2). However, it is likely a combination of these mechanisms that drives the coevolution of S*. ‘fitti’* strains within Caribbean acroporids. In either case, *S. ‘fitti’* genetic diversity is tied to that of its endangered hosts.

### Host selectivity of symbiont strains in a horizontal symbiont transmission

Broadcast spawning coral species acquire symbiotic partners horizontally each generation during the aposymbiotic larval stage (Baird, Guest, & Willis 2009). Thus, it is perplexing that coral lineages remain specific to a single symbiont strain, despite the potential to select a novel symbiont that might expand its physiological capacity and range of suitable habitats (Barneah, Weis, Perez, & Benayahu 2004; Chan, Lewis, Neely, & Baums 2019). Though initial intra-family, intra-genus, and intra-specific symbiont diversity is observed during early months of development, the diversity wanes and reverts to the dominant (adult) symbiont species shortly thereafter (Abrego et al. 2009; Poland & Coffroth 2017). Specificity between adult broadcast spawning corals and their symbionts is commonly observed at the species level with multi-marker and microsatellite approaches (Chan et al. 2019; LaJeunesse 2001; Lewis et al. 2019). Here, we add population genomic analyses using SNP data to reveal that sub-species level partitioning occurs between *S. ‘fitti’* and its three Caribbean acroporid hosts (Figs. 2–4). Though adult colonies of Caribbean acroporids primarily associate with *S. ‘fitti’,* they can occasionally harbor other genera of Symbiodiniaceae (*Breviolum* spp., *Durusdinium trenchii* and *Cladocopium* spp.), but these associations are often transitory and revert to *S. ‘fitti’* over time (Baums, Johnson, Devlin-Durante, & Miller 2010; Thornhill, LaJeunesse, Kemp, Fitt, & Schmidt 2006b).

The degree of selectivity of the cnidarian host when accepting a symbiotic partner likely has a role in maintaining the long-term specificity in these mutualisms. Host selectivity is in part modulated by cell recognition pathways such as lectin-glycan interactions and might contribute to the coevolution of *S. ‘fitti’* and its respective host acroporids (Davy, Allemand, & Weis 2012; Logan, LaFlamme, Weis, & Davy 2010; Parkinson et al. 2018; Weis, Reynolds, deBoer, & Krupp 2001; Wood-Charlson, Hollingsworth, Krupp, & Weis 2006). Though the *exact* role of cell-signaling in host selectivity has yet to be fully described, the specificity between partners is maintained by specialization pressures (LaJeunesse et al. 2018; Parkinson et al. 2018; Wood-Charlson et al. 2006). The ~160 million years following the widespread adaptive radiation of stony corals and Symbiodiniaceae has resulted in their current inter-dependence on one another (LaJeunesse et al. 2018). Consequently, the various symbiotic pairings remain adapted to meet the unique biochemical and metabolic demands of each host microenvironment (Barott, Venn, Perez, Tambutté, & Tresguerres 2015; Sogin, Anderson, Williams, Chen, & Gates 2014). The different morphologies and corallite structures possessed by each acroporid results in different light scattering properties and ultimately, light availability to the resident endosymbiont (Enríquez et al. 2017). These differences likely lead to strong selection pressures unique to each acroporid taxa. Ultimately, the specificity between *S. ‘fitti’* strains and their acroporid hosts could be maintained by its ability (or lack thereof) to meet the metabolic needs of its host and adapt to the microenvironment created by its host.

### *The role of symbiont selectivity as a driver of* S. ‘fitti’ *intraspecific genomic variation*

Akin to role of host selectivity, that of the symbiont is also important for upholding partner specificity in symbioses. The diversity and abundance of Symbiodiniaceae in the water column is not necessarily proportional to their counterparts partaking in mutualisms with reef-building corals (Cunning, Yost, Guarinello, Putnam, & Gates 2016; Littman, van Oppen, & Willis 2008; Manning & Gates 2008). However, some lineages of Symbiodiniaceae possess the ability to establish symbiosis with non-coral invertebrates (Cunning et al. 2016; Decelle et al. 2018). The total diversity of *S. ‘fitti’* likely spans beyond strains that inhabit acroporids and may incorporate free-living conspecifics living in the water column, sediments, etc. Therefore, the slight differences in allelic composition of each *S. ‘fitti’* strain may be a result of the differential host preference of the available symbionts (Figs. 2–4). However, symbiotic *S. ‘fitti’* strains might be attracted to the different microbial composition and abundance in the water column adjacent to a coral colony that constitute the ‘ecosphere’ surrounding each acroporid (Weber, González-Díaz, Armenteros, & Apprill 2019). Similarly, the food associated with each ‘ecospheres’ may attract different Symbiodiniaceae (Pollock et al. 2018; Weber et al. 2019). Putative intraspecific variation in the swimming availability and chemosensory responses of *S. ‘fitti’* may also, in part, dictate which Symbiodiniaceae persist in each ecosphere (Fitt 1984; Fitt 1985; Fitt, Chang, & Trench 1981; Kamykowski, Reed, & Kirkpatrick 1992). Future experimental validation of intraspecific variation in *S. ‘fitti’* swimming and chemosensory ability and how it pertains to selectivity of their acroporid hosts (and ‘ecospheres’) will shed light on how inter-partner specificity is maintained.

### *Coevolution as a driver of* S. ‘fitti’ *intraspecific genomic variation*

Coevolution is the process by which two interacting species reciprocally adapt to each other (sensu Janzen 1980). The coevolution of cnidarian-dinoflagellate mutualisms, in part, has resulted in the long-term fidelity between partners (Figs. 2–4; LaJeunesse et al. 2018; Stanley 2006). Though the fossil record supports the coevolution of these mutualisms (Muscatine et al. 2005; Stanley 2006), experimental follow up and verification has received far less attention. Advances in phylogenomics reveal extensive differentiation of genomic features and gene family enrichment when comparing symbiotic and free-living *Symbiodinium* spp., a potential biproduct of coevolving with their hosts (González-Pech, Bhattacharya, Ragan, & Chan 2019a; González-Pech, Ragan, & Chan 2017; González-Pech et al. 2019b). Similarly, population genetic analyses of host and symbiont reveals widespread long-term partnerships between several cnidarian species and their endosymbiotic dinoflagellates, which might be the biproduct of coevolution (Baums et al. 2014; O’Donnell et al. 2018; Poland & Coffroth 2017; Thornhill et al. 2006a; Thornhill et al. 2017; Thornhill et al. 2006b). Therefore, the correspondence of the allelic variation of *S. ‘fitti’* to its host acroporids is likely, in part, the result of coevolution. The STAGdb genotyping array (SNPchip) can be harnessed to experimentally verify the contributions of coevolution, host specificity, and symbiont selectivity to the evolutionary dynamics of *Acropora-S. ‘fitti’* symbioses’ (Kitchen et al. 2020).

### The special case of the F1 coral hybrid as habitat

The shared history (~2.6-3.6 million years of coexistence) between parents *A. palmata* and *A. cervicornis* with their symbiont, *S. ‘fitti’* may have allowed sufficient time for co-evolutionary processes to play out (and so may help explain the strain differentiation of *S. ‘fitti’* by host; Figs 2–4; Budd & Johnson 1999; McNeill et al. 1997). However, the situation differs for their first-generation hybrid, *A. prolifera*, which cannot directly respond to selection pressure from *S. ‘fitti’* via differential successful sexual reproduction of its colonies because most are sterile (Vollmer & Palumbi 2002). Thus, any changes in the allele frequencies of *A. prolifera* are restricted to somatic mutations occurring within their lifetime which can be on the order of hundreds of years (Irwin et al. 2017). Though the fossil record of *A. prolifera* is rather sparse (McNeill et al. 1997), *S. ‘fitti’* may have encountered these colonies over many thousands of years as they are generated anew with each hybridization event. Thus, while a host co-adaptive response is unlikely, *S. ‘fitti’* may have evolved strains that preferentially colonize *A. prolifera*.

### Environmental differentiation as a driver of intraspecific genomic variation

Environmental differences create variation in partner selectivity (Thrall, Hochberg, Burdon, & Bever 2007). *S. ‘fitti’* population dynamics are confounded with differences in host habitat preferences including light, temperature, nutrient concentration, and food availability in the water column (Crossland & Barnes 1983; Miller 1995; Terraneo et al. 2019; Williams et al. 2018). The inverse relationship between depth and light availability is a common driver of coral and Symbiodiniaceae zonation (Bongaerts et al. 2015a; Bongaerts et al. 2015b; Bongaerts et al. 2017; Fogarty 2012; Goulet, Lucas, & Schizas 2019; LaJeunesse 2002; Serrano et al. 2014; Serrano et al. 2016). Throughout much of their distribution, Caribbean acroporid species reside in different habitats (Fogarty 2012; Goreau 1959). Specifically, *A. cervicornis* occupies a lower light habitat (~10m depth) relative to its high-light dwelling (~3m depth) sibling species *A. palmata* (Fogarty 2012; Goreau 1959; LaJeunesse 2002). Although the hybrid’s depth distribution often overlaps with *A. palmata,* it can also be found in less than 1m of water (Fogarty 2012). Therefore, adaptation to different light availabilities may correspond to genomic differentiation between the shallow *A. palmata*-*S. ‘fitti’* and *A. prolifera*-*S. ‘fitti’* versus deep *A. cervicornis-S. ‘fitti’* (Figs. 2–4; Finney et al. 2010; Kirk, Andras, Harvell, Santos, & Coffroth 2009). Microenvironments created by light attenuation at depth may lead to range-limited dispersal of *S. ‘fitti’* and modulate the available pool of symbionts (Finney et al. 2010; Serrano et al. 2016). Furthermore, the differences in the skeletal morphology of acroporids result in different light scattering properties that likely have profound effects on the light availability and microenvironments for their resident *S. ‘fitti’* (Enríquez et al. 2017; Gladfelter 1983; Gladfelter 2007). The different light regimes, skeletal features and flow fields associated with each acroporid may exert some selection pressure on symbiont strains.

### *Genomic basis for extended phenotypes in* S. ‘fitti’ *- acroporid symbiosis*

The physiological capacity of the holobiont (coral and symbiont) hinges upon the specific partner pairings as well as external (environmental) drivers (Parkinson & Baums 2014). The large number of non-synonymous SNPs differentiating the *S. ‘fitti’* strains among their hosts may demarcate the onset of eventual speciation (Fig. 5; Tables S7-10) although it is difficult to know what barriers to gene flow may exist that allow for such a process. Changes in amino acid sequences and protein efficiency resulting from these mutations may serve as the genomic basis causing intraspecific variation in physiological aptitude (Parkinson et al. 2015a; Parkinson & Baums 2014). Similarly, the candidate genes under selection identified by BayeScan and PCAdapt may underlie the strain differentiation by host species (Tables S7-10). Non-synonymous mutations in putative cytosolic oligopeptidase and alpha-agarase regions may result in the subtle alteration of zinc and calcium ion binding, respectively, which in turn likely contribute to variation in the physiological capacity of *S. ‘fitti’* (Table S7-10; Kmiec, Teixeira, Murcha, & Glaser 2016; Zhang et al. 2018). Further, these genotypic and phenotypic differences may facilitate the adaptation of each *S. ‘fitti’* strain to the internal and external microenvironments associated with each host niche and meeting their metabolic and nutritional demands (Figs. 2–5; Tables S7-10; Hemond, Kaluziak, & Vollmer 2014; Muscatine, Porter, & Kaplan 1989; Reich, Rodriguez, LaJeunesse, & Ho 2020; Sogin et al. 2014).

### Conclusion

We show here that the population genetic structure of *Symbiodinium ‘fitti*’ is, in part, explained by its host association. Because the host species occupy different habitats, we cannot yet disentangle the role of host versus depth as a potential driver of population genetic structure. However, the genomic resources for the *S. ‘fitti’* - acroporid system described here can be used in future studies to determine whether, and to what degree, the observed variation of allelic composition is a result of host selectivity, symbiont selectivity, coevolution, environmental differentiation, or a combination of these mechanisms. The appreciation of population genetic structure and evolutionary dynamics of both coral holobiont partners will better inform the genomic underpinnings of their phenotypes and physiological capacity.

## Acknowledgements

We thank the PSU genomics facility for assistance with library prep and sequencing. We thank Prof. Todd LaJeunesse for assistance with the *psbA* phylogeny. Funding for this project was supported by NSF-OCE-1537959 (to IBB) and NSF-OCE-1538469 (to NDF). HGR was supported through NSF-OCE-1636022 (to T. LaJeunesse). Permits for samples include Florida: CRF permit numbers CRF-2017-009, CRF-2017-012, NOAA FKNMS permit numbers FKNMS-2011-159-A4, FKNMS-2001-009, FKNMS-2014-148-A2, and FKNMS-2010-130-A, Belize: CITES Permit 0385, 7487 and 7488; Curacao: CITES Permit 16US784243/9 and 12US784243/9; and USVI Department of planning and natural resources, Division of fish and wildlife DFW14017T.

## Data availability

Sequences are under NCBI SRA PRJNA473816. Code for data analysis and figure generation is available on github (https://github.com/hgreich/Sfitti).

## Author contributions

Conceived the project: SAK, IBB, HGR. Obtained funding: IBB, NDF. Mentorship: IBB, SAK. Field collections of corals: SAK, IBB, NDF. Molecular work: SAK, MDD. Bioinformatic analyses: HGR, SAK, KHS. Wrote the paper: HGR, SAK, IBB, NDF, KHS.

## Tables

Table S1: Sample information including host taxa, location, sampling depth, sequencing platform, and accession number. Per sample information on read counts, mapping rates, and SNP summary statistics are also included. Multi-locus genotypes from 12 microsatellite loci, 58,538 “high quality” SNPs, and 6,813 “conservative” SNPs with no missing data are included.

Table S2: Genome assembly summary statistics for *Symbiodinium ‘fitti’* sample PFL14120.

Table S3: Annotation information for the 24,000+ genes in *Symbiodinium ‘fitti.’*

Table S4: Annotation information and selection outlier statistics for *Symbiodinium ‘fitti’* SNPS (12,700) that are predicted to fall in coding regions.

Table S5: AMOVA of indicates that ~12% of *Symbiodinium ‘fitti’* variation is due to host taxon whereas negative % variation is due to location.

Table S6: Summary statistics from Structure Harvester including the Evanno’s *delta* K method which predicted three main clusters.

Table S7: Selection outlier statistics and per SNP summary statistics loci identified as selection outliers by BayeScan (370) and PCAdapt (4,987). 307 selection outlier SNPs were shared between the two programs.

Table S8: Summary statistics for the predicted downstream effects of SNPs (generated by the SNPeff program). 60,373 (84.43%) modifier/non-coding, 3,629 (5.08%) moderate/mostly harmless, 7,451 (10.42%) low impact variants that might change protein efficiency/effectiveness, and 51 (0.07%) highly disruptive SNPs were identified.

Table S9: Predicted downstream effects (SnpEff) and annotation information for variants in coding regions.

Table S10: Predicted downstream effects (SnpEff) and annotation information for 12 variants in coding regions that were identified as selection outliers by both PCAdapt and BayeScan.

---

**Fig. S1:**
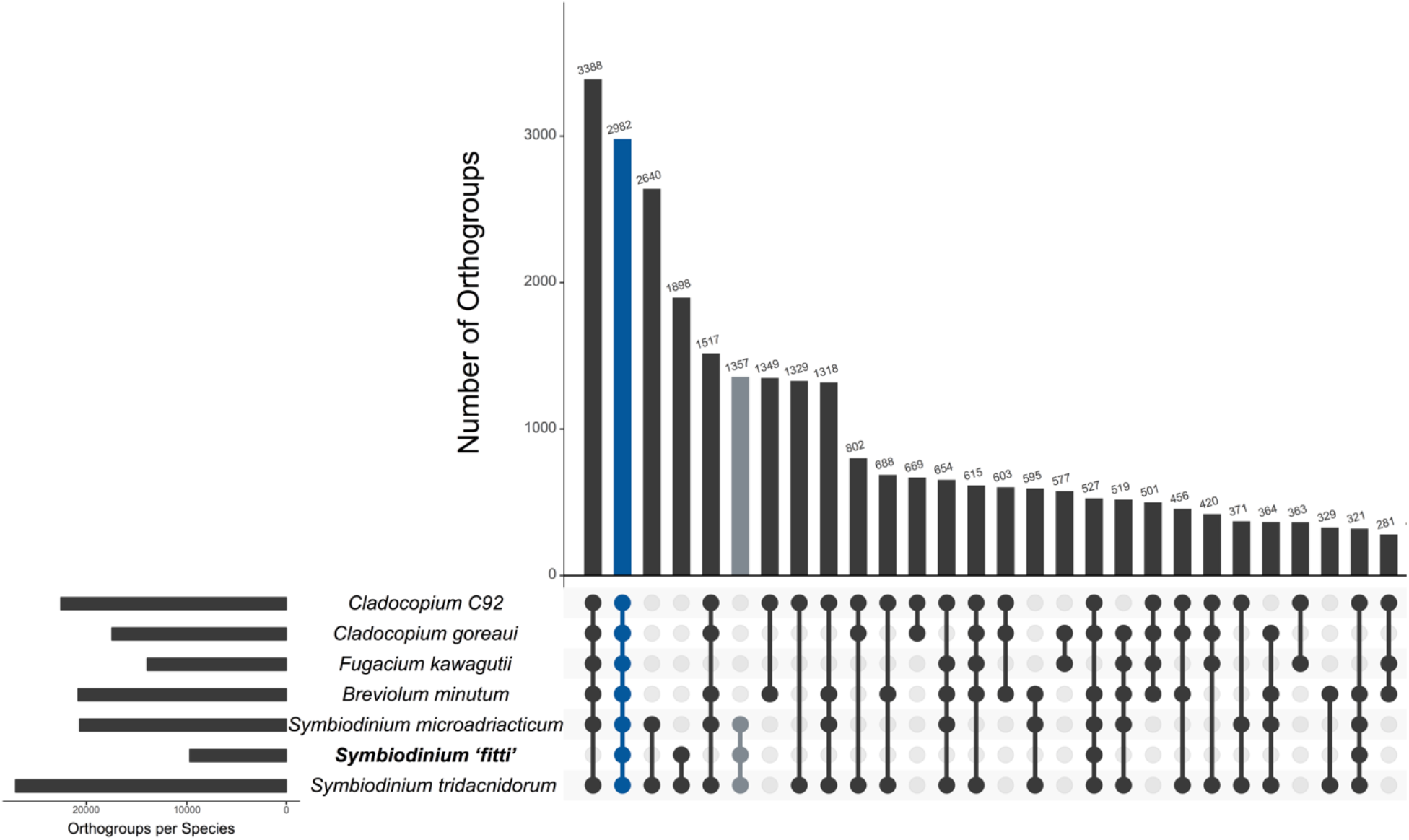
Gene families shared between different Symbiodiniaceae lineages (including *Symbiodinium ‘fitti’*).

**Fig. S2:**
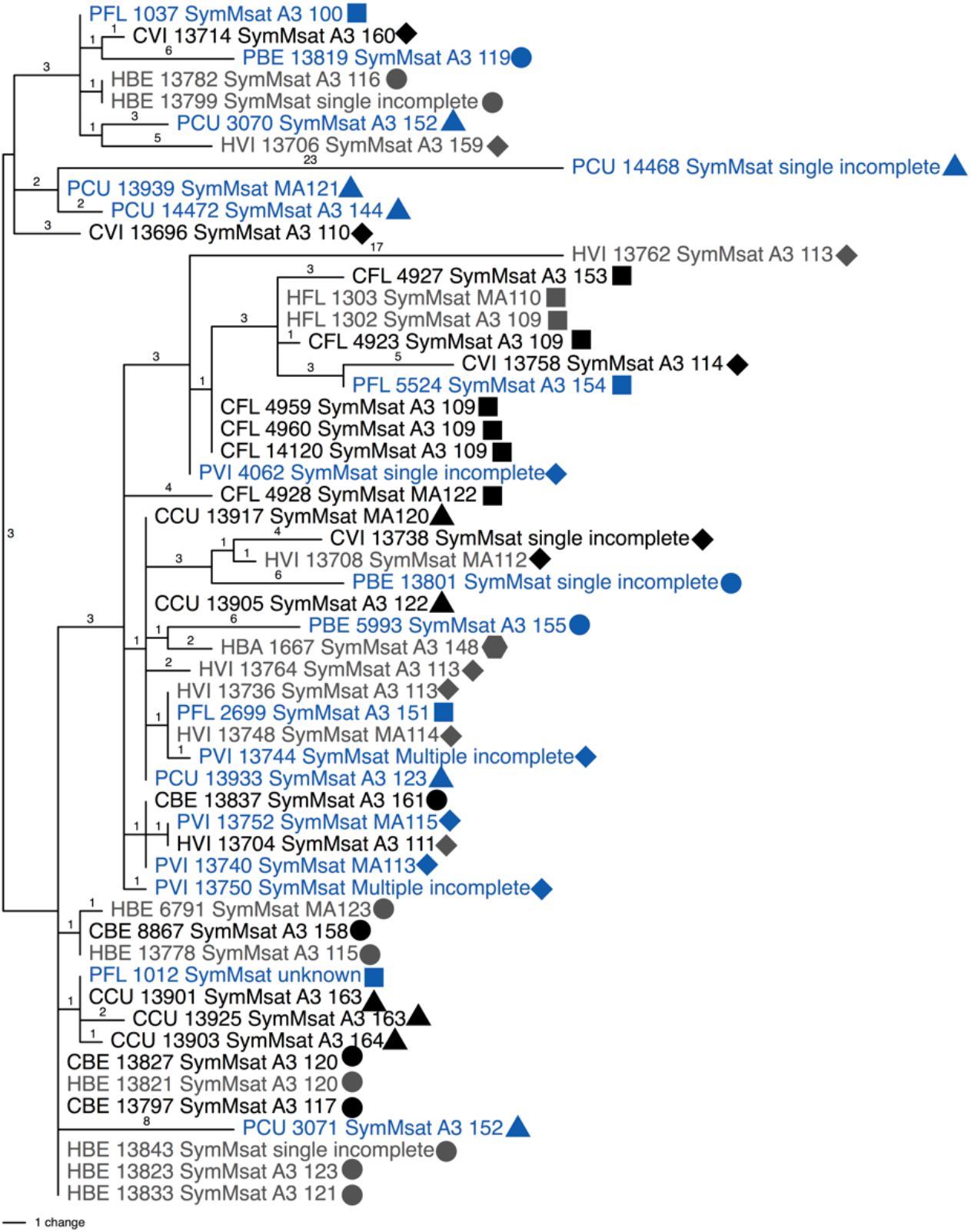
Intragenomic variation of *Symbiodinium ‘fitti’* is at the sub-species level. Phylogeny of the *psbA* non-coding region, a commonly used marker used to help delimit *Symbiodinium* species indicates that *S. ‘fitti’* in its three different hosts are all one species.

**Fig. S3:**
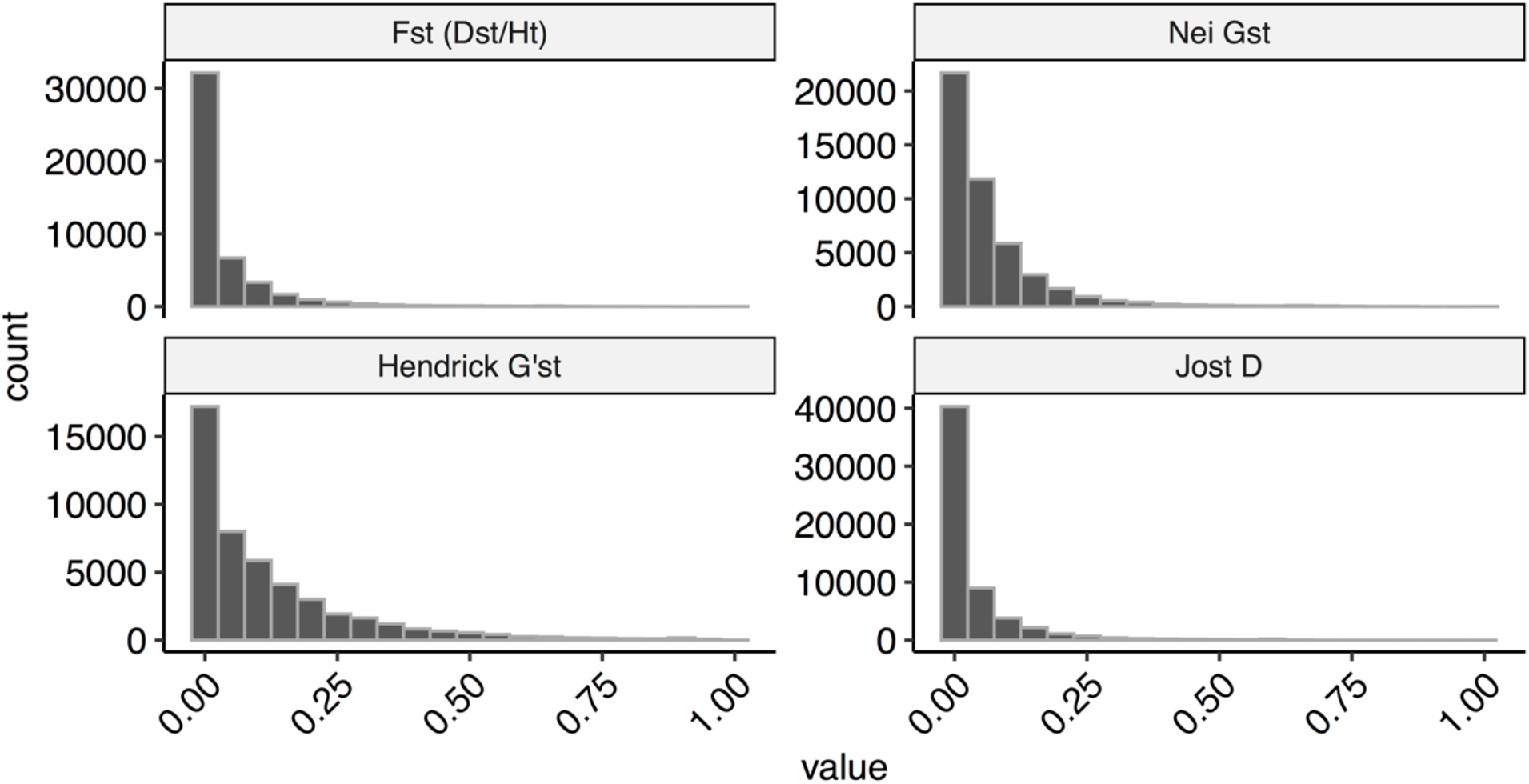
Histogram of per SNP fixation levels for 58,538 genotyping SNPs.

